# Genome-wide association study of 40,000 individuals identifies two novel loci associated with bipolar disorder

**DOI:** 10.1101/044412

**Authors:** Liping Hou, Sarah E. Bergen, Nirmala Akula, Jie Song, Christina M. Hultman, Mikael Landén, Mazda Adli, Martin Alda, Raffaella Ardau, Bárbara Arias, Jean-Michel Aubry, Lena Backlund, Judith A. Badner, Thomas B. Barrett, Michael Bauer, Bernhard T. Baune, Frank Bellivier, Antonio Benabarre, Susanne Bengesser, Wade H. Berrettini, Abesh Kumar Bhattacharjee, Joanna M. Biernacka, Armin Birner, Cinnamon S. Bloss, Clara Brichant-Petitjean, Elise T. Bui, William Byerley, Pablo Cervantes, Caterina Chillotti, Sven Cichon, Francesc Colom, William Coryell, David W. Craig, Cristiana Cruceanu, Piotr M. Czerski, Tony Davis, Alexandre Dayer, Franziska Degenhardt, Maria Del Zompo, J. Raymond DePaulo, Howard J. Edenberg, Bruno Étain, Peter Falkai, Tatiana Foroud, Andreas J. Forstner, Louise Frisén, Mark A. Frye, Janice M. Fullerton, Sébastien Gard, Julie S. Garnham, Elliot S. Gershon, Fernando S. Goes, Tiffany A. Greenwood, Maria Grigoroiu-Serbanescu, Joanna Hauser, Urs Heilbronner, Stefanie Heilmann-Heimbach, Stefan Herms, Maria Hipolito, Shashi Hitturlingappa, Per Hoffmann, Andrea Hofmann, Stephane Jamain, Esther Jiménez, Jean-Pierre Kahn, Layla Kassem, John R. Kelsoe, Sarah Kittel-Schneider, Sebastian Kliwicki, Daniel L. Koller, Barbara König, Nina Lackner, Gonzalo Laje, Maren Lang, Catharina Lavebratt, William B. Lawson, Marion Leboyer, Susan G. Leckband, Chunyu Liu, Anna Maaser, Pamela B. Mahon, Wolfgang Maier, Mario Maj, Mirko Manchia, Lina Martinsson, Michael J. McCarthy, Susan L. McElroy, Melvin G. McInnis, Rebecca McKinney, Philip B. Mitchell, Marina Mitjans, Francis M. Mondimore, Palmiero Monteleone, Thomas W. Mühleisen, Caroline M. Nievergelt, Markus M. Nöthen, Tomas Novák, John I. Nurnberger, Evaristus A. Nwulia, Urban Ösby, Andrea Pfennig, James B. Potash, Peter Propping, Andreas Reif, Eva Reininghaus, John Rice, Marcella Rietschel, Guy A. Rouleau, Janusz K. Rybakowski, Martin Schalling, William A. Scheftner, Peter R. Schofield, Nicholas J. Schork, Thomas G. Schulze, Johannes Schumacher, Barbara W. Schweizer, Giovanni Severino, Tatyana Shekhtman, Paul D. Shilling, Christian Simhandl, Claire M. Slaney, Erin N. Smith, Alessio Squassina, Thomas Stamm, Pavla Stopkova, Fabian Streit, Jana Strohmaier, Szabolcs Szelinger, Sarah K. Tighe, Alfonso Tortorella, Gustavo Turecki, Eduard Vieta, Julia Volkert, Stephanie H. Witt, Adam Wright, Peter P. Zandi, Peng Zhang, Sebastian Zollner, Francis J. McMahon

## Abstract

Bipolar disorder (BD) is a genetically complex mental illness characterized by severe oscillations of mood and behavior. Genome-wide association studies (GWAS) have identified several risk loci that together account for a small portion of the heritability. To identify additional risk loci, we performed a two-stage meta-analysis of >9 million genetic variants in 9,784 bipolar disorder patients and 30,471 controls, the largest GWAS of BD to date. In this study, to increase power we used ~2,000 lithium-treated cases with a long-term diagnosis of BD from the Consortium on Lithium Genetics, excess controls, and analytic methods optimized for markers on the X-chromosome. In addition to four known loci, results revealed genome-wide significant associations at two novel loci: an intergenic region on 9p21.3 (rs12553324, *p* = 5.87×10^−9^; odds ratio = 1.12) and markers within *ERBB2* (rs2517959, *p* = 4.53×10^−9^; odds ratio = 1.13). No significant X-chromosome associations were detected and X-linked markers explained very little BD heritability. The results add to a growing list of common autosomal variants involved in BD and illustrate the power of comparing well-characterized cases to an excess of controls in GWAS.

## INTRODUCTION

Bipolar disorder (BD) is a common, chronic and an episodic mental disorder characterized by disruptive oscillations of mood and behaviour. The lifetime prevalence estimate of the US population is about 2% (BD-I and BD-II), but exceeds 2% for sub-threshold conditions (1, 2). The elevated morbidity and mortality associated with BD make it a major public health problem. Despite advances in recent years, the underlying neurobiology of BD remains largely unknown.

The overall heritability of BD has been estimated to be more than 70% based on twin studies (3, 4). Genome-wide association studies (GWAS) have identified several risk loci. These include markers near *ADCY2, ANK3, CACNA1C, TENM4, SYNE1, TRANK1*, and a tight cluster of genes on chromosome 3p21, among others (5–9). These loci account for only a small portion of the heritability of BD, suggesting that additional risk loci remain to be discovered.

The highly polygenic architecture of BD (10) suggests that identification of additional risk loci will require larger samples than have been studied so far. As the diagnosis of BD can be challenging, great care must be taken in the selection of cases (11). Accordingly, the ascertainment of well-characterized cases has proven to be a limiting factor. To address this problem, the present study augmented previously published case sets with a large set of well-characterized cases followed on lithium for at least 6 months and assembled by the Consortium on Lithium Genetics (ConLiGen) (12, 13). These cases were included in a recently published GWAS of lithium response (13), but have heretofore not been used for GWAS of BD itself. Since an excess of controls beyond the traditional 1:1 case:control ratio can confer a meaningful increase in power in GWAS (14–16), we have also included over twenty thousand genotyped controls drawn from public databases. Most have not, to our knowledge, been included in previous GWAS of BD.

Surprisingly few of the published GWAS of BD have reported results for X chromosome markers, even though family and genetic linkage studies have long suggested a role for X-linked genes in BD (17–19). While the smaller effective sample size of X-linked markers necessarily leads to reduced power relative to autosomal markers (20), omission of the X-chromosome represents a considerable loss of potential association signals, since it comprises approximately 5% of the female and 2.5% of the male genome. One reason for the omission may be the relative paucity of association methods that correctly account for the added complexities of X-linked markers. Recent advances have improved the available analytic tools (21, 22), and we employ one such tool in the present study. We also employ a large X-chromosome imputation reference panel from the 1000 Genomes Project (23) that was not available during the first generation of GWAS.

In summary, the present study aimed to identify additional BD risk loci by carrying out a GWAS with new cases, excess controls, and analytic methods optimized for the X-chromosome. The most significant single nucleotide polymorphisms (SNPs) were tested for association in an independent replication sample of about 2,300 cases and 3,500 controls from two independent GWAS of BD. While we did not detect any genome-wide significant variants on chromosome X, we did find genome-wide significant evidence for common risk variants at two novel and four known autosomal loci. The results add to a growing list of common autosomal markers associated with BD and illustrate the power of well-characterized cases, combined with an excess of controls, to identify previously unknown loci involved in common, polygenic disorders.

## RESULTS

A total of 7,647 cases and 27,303 controls were analyzed in Stage 1 (Table 1), in which a total of 9,692,718 autosomal markers passed quality control. The Stage 1 studies had >90% power to detect association at the significance level of *P* <1 x 10^−6^ with a common autosomal allele that confers a genotype-relative risk (GRR) of 1.15. The p-value distributions were unbiased for each of the sub-studies: all standardized genome-wide inflation factors (λ_1000_) were ≤1.07 (Figure S1). Meta-analysis of the Stage 1 studies identified 62 variants that exceeded the standard genome-wide significance threshold (Table S1). All lay within two known risk loci^7,9^, one SNP (rs9834970, *P* = 3.19 × 10^−8^, OR = 0.88) lay near the gene *TRANK1*; all others were located in the gene *MAD1L1*. All 179 variants with fixed-effect *P* < 10^−6^ were carried forward to the Stage 2 samples for further validation (Table S1). After LD-pruning at r^2^= 0.2, the Stage 1 results appeared to represent 14 distinct regions.

**Table 1.** Study sample characteristics

| Study | Female (%) | Platform | Samples | Cases | Controls | Total |
| --- | --- | --- | --- | --- | --- | --- |
| Stage 1 |  |  |  |  |  |  |
| Study-1 | 49.2 | Affymetrix 6.0 | GAIN-NIMH | 974 | 1,032 | 2,006 |
| Study-2 | 54.7 | Affymetrix 500K | WTCCC1-BD | 1,844 | 2,932 | 4,776 |
|  |  |  | STEP-BD | 945 | 650 | 1,595 |
| Study-3 | 52.2 | Affymetrix 6.0 | TGEN1 | 1,184 | 400 | 1,584 |
|  |  |  | ConLiGen | 32 | 0 | 32 |
|  |  |  | WTCCC2 | 0 | 2724 | 2,724 |
| Study-4 | 48.7 | HumanHap550 | BoMA | 628 | 1,291 | 1,919 |
|  |  |  | dbGaP<sup>a</sup> | 0 | 1,462 | 1,462 |
| Study-5 | 53.7 | HumanOmni2.5M | ConLiGen | 416 | 0 | 416 |
|  |  |  | dbGaP<sup>a</sup> | 0 | 2,741 | 2,741 |
| Study-6 | 50.0 | Illumina610/660/1M/OE | ConLiGen | 1,624 | 0 | 1,624 |
|  |  |  | dbGaP<sup>a</sup> | 0 | 14,071 | 14,071 |
| Stage 2 |  |  |  |  |  |  |
| Swedish sample 1 | 52.2 | Affymetrix 6.0 | sw34 | 796 | 1,986 | 2,782 |
| Swedish sample 2 | 57.6 | OmniExpress | sw6 | 1,341 | 1,182 | 2,523 |
| Grand total | 51.7 |  |  | 9,784 | 30,471 | 40,255 |

<sup>a</sup> Details on contributing samples from dbGaP are given in Table S4.

Meta-analysis of the combined Stage 1 and Stage 2 studies, comprising 9,784 cases and 30,471 controls, had >90% power to detect association at the genome-wide significance level of *P* <5.0 x 10^−8^ with a common autosomal allele that confers a GRR of 1.15. The meta-analysis produced an unbiased distribution of p-values (Figures S2): The genome-wide inflation factor was 1.08, while the standardized genomic inflation factor (λ_1000_) was ~1.01.

Six autosomal loci exceeded genome-wide significance (Figure 1). Four of these loci have been described before (Figure S3) (7–9, 24). The remaining two loci are novel BD risk loci (Figure 2). None of the six genome-wide significant loci identified here demonstrated significant heterogeneity in effect sizes across the samples studied (Table 2).

**Figure 1.**
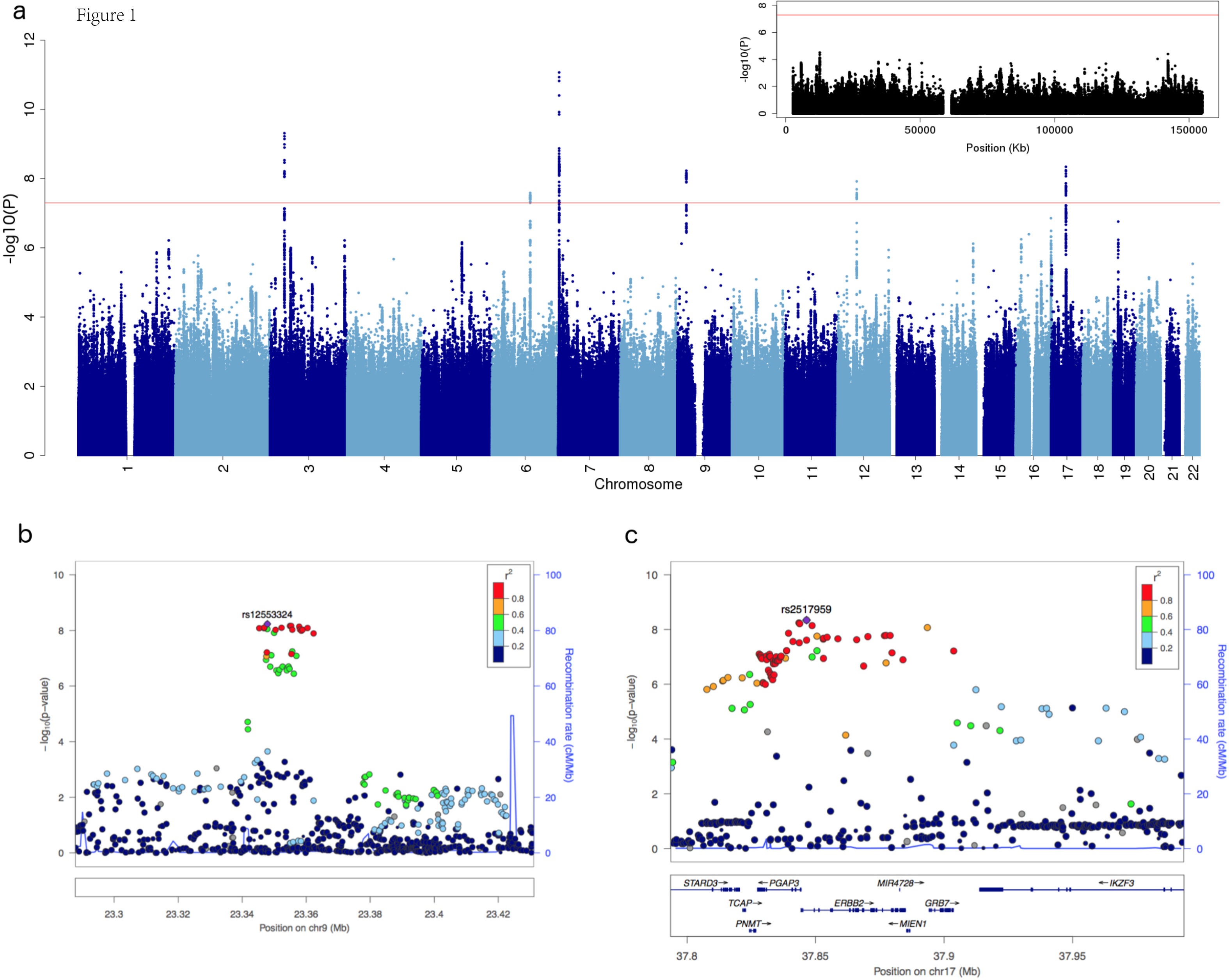
**Manhattan plot and regional association plots.** (a) Manhattan plot for all analyzed markers (the inset is for markers on the X chromosome); (b,c) regional association plots for the two novel risk loci (left: 9p21.3, right: *ERBB2* region).

**Figure 2.**
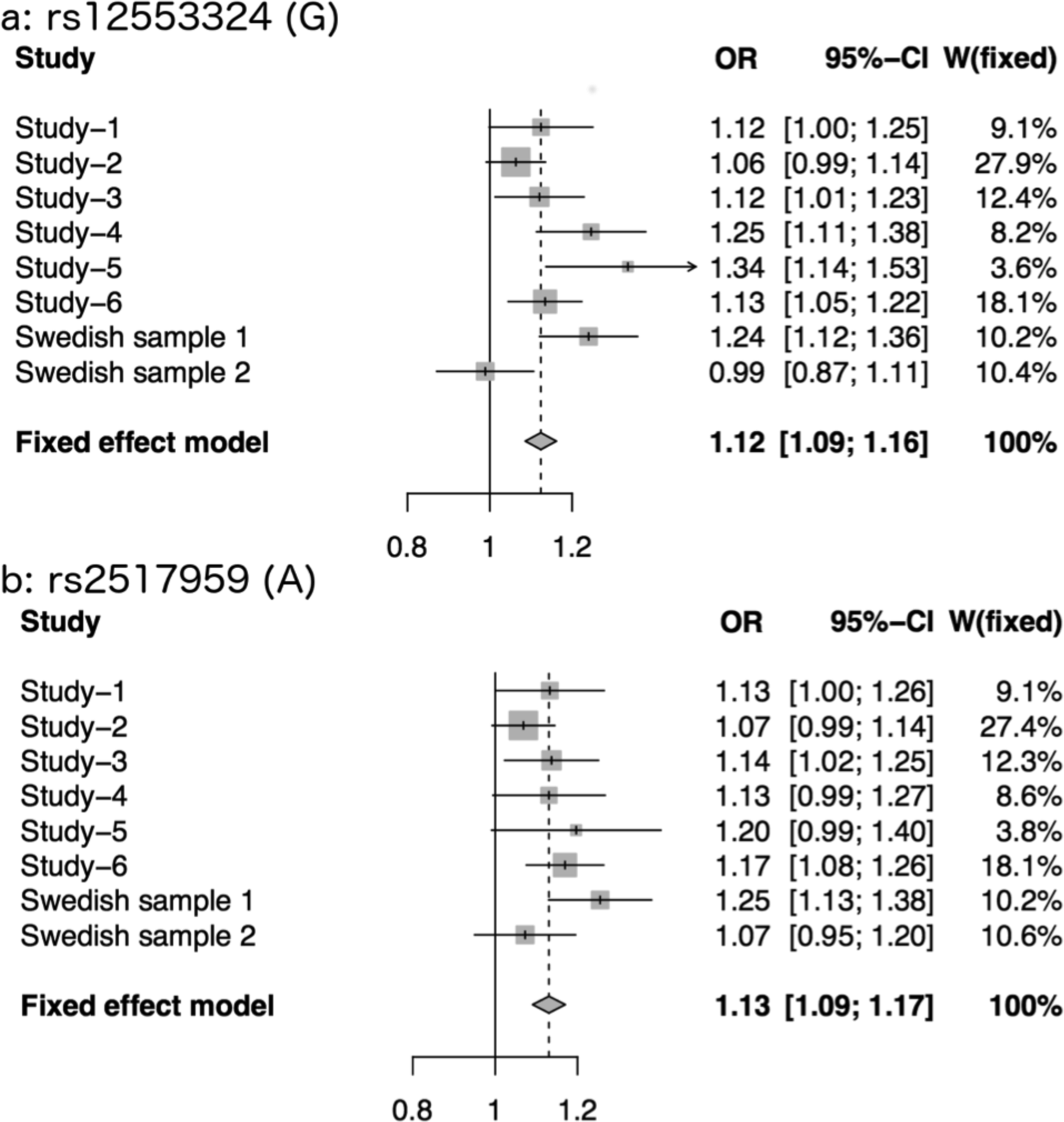
**Genetic effect sizes for the novel risk loci.** Forest plots displaying the odds ratios
(OR) and 95% confidence intervals for the most significant SNPs in the (a) 9p21.3 region and (b) the *ERBB2* region. Horizontal lines indicate the 95% confidence interval of the OR for each study, with a shaded box around the point estimate, drawn proportional to the sample size. The diamond indicates the overall weighted OR for all samples included in the meta-analysis.

**Table 2.**
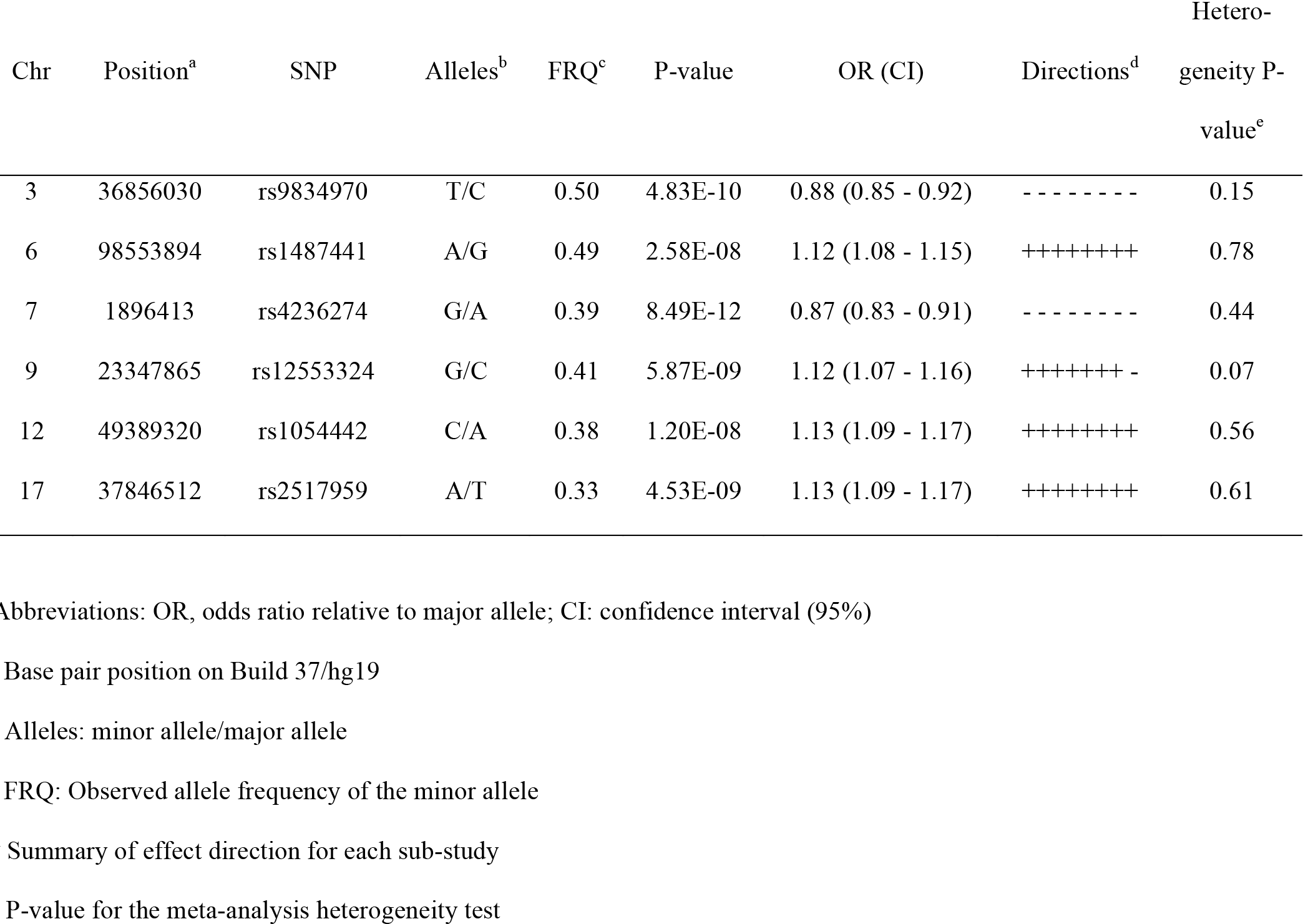
Top SNPs in regions showing genome-wide significant evidence for association with BD in the combined analysis

The first novel locus is located on chromosome 17q12. The most significant SNP (rs2517959, *P* = 4.53 × 10^−9^, OR = 1.13) is located in an intronic region of the gene, *ERBB2*, which encodes a receptor tyrosine kinase. Several other genes also lie nearby (Figure 1).

The top SNP in the second novel locus, rs12553324, lies within an intergenic region on chromosome 9p21.3 (*P* = 5.87 × 10^−9^, OR = 1.12). One SNP in moderate LD (rs10965780; r^2^ = 0.604 (25)) lies within the promoter flanking region of *ELAVL2*, which encodes a neuron-specific RNA binding protein (Refseq, November 2015).

The most significant association signal in this study falls within the *MAD1L1* gene on chromosome 7p22.3, and has been reported by previous studies of BD or BD plus schizophrenia (SCZ) (8,26). The top SNP, rs4236274, is located in an intron of *MAD1L1* (*P* = 8.49 × 10^−12^, OR = 0.87). An additional 60 variants at this locus surpassed the genome-wide significance threshold.

The second most significant finding in this study lies near the gene *TRANK1* on chromosome 3p22.2. This has been identified as a genome-wide significant risk locus for BD by two previous GWAS with partially overlapping samples (7, 9). The same SNP and allele of rs9834970 from those two studies was also significantly associated with BD in the present study (*P* = 4.83 × 10′ ^10^, OR = 0.88).

Twelve variants on chromosome 6q16.1 and ten variants on 12q13.1 also met the genome-wide significance threshold. The top SNP within the 6q16.1 locus (rs1487441, *P* = 2.58 × 10^−8^, OR = 1.12) is located in an intergenic region between *MIR2113* and *POU3F2*. Within the 12q13.1 locus, the top SNP (rs1054442, *P* = 1.20 × 10^−8^, OR = 1.13) is located within the 3’–UTR of *DDN*, which encodes dendrin, a cytoskeletal protein expressed at the synapse (27).

We also note nominally significant (*P* <0.01) support for most of the loci identified in previous GWAS of BD, including markers on chromosome 3p21, and near the genes *ADCY2, ANK3, CACNA1C, LMAN2L, NCAN, TENM4*, and *SYNE1* (Table S2).

Several of the identified loci contained multiple SNPs in tight LD. To clarify whether each locus represented a single association signal, we conducted an approximate conditional analysis using GCTA (see Methods). The results suggested that each of the six GWAS-significant loci (Table 2) was consistent with only one distinct signal (Table S3).

In the X-chromosome analysis, a total of 218,707 markers passed stringent quality control. The power analysis suggested that the Stage 1 studies had 65% power to detect a common X-linked allele that confers a GRR of 1.2 at the significance level of p<1 x 10^−6^, while the combined Stage 1 and 2 meta-analysis had 66% power to detect association with the same allele at the genome-wide significance level of p<5.0 x 10^−8^. No X-chromosome markers met the p<1 x 10^−6^ threshold to be carried forward from Stage 1 to Stage 2. Consequently we did not identify any genome-wide significant signals on the X chromosome (Figure 1).

We also assessed the relative distribution of genomic heritability represented in the Stage 1 studies. Consistent with a highly polygenic architecture, there was a strong linear relationship between the genomic heritability attributable to each chromosome and chromosome length (*P* = 0.0004, R^2^ = 0.45) (Figure S4), except for the X-chromosome. Unlike autosomal SNPs, X-linked SNPs explained an unexpectedly small proportion of the genomic heritability of BD in this study (0.2%, Figure S4).

## DISCUSSION

This study is the first GWAS of BD to include samples from the ConLiGen study (13) and to our knowledge the largest published to date. The full set of summary results is available for download at http://www.nimh.nih.gov/hgb-data/BPGWASmetaresults.tar.gz. The analysis identified two novel and four known BD risk loci. The results also provided nominally significant support for most loci identified in previous GWAS of BD. No significant X-chromosome associations were detected and X-linked markers explained very little of the genomic heritability of BD.

This study has several limitations. The total sample size is still too small to identify risk variants with very small effect sizes or low frequencies, especially any residing on the X chromosome. Larger scale studies, such as those ongoing within the Psychiatric Genomics Consortium, will be needed to identify such loci. It has been shown that meaningful increases in statistical power for case-control studies can be obtained by increasing the control-to-case ratio in the range of 4 to 5 (14, 15). Thus we used an excess of controls, including unscreened controls from WTCCC2 and dbGaP. Such unscreened controls are generally easy to obtain and inexpensive, but the actual gain in power may be less than the sample size alone suggests, since some might qualify as cases if examined. However, the population prevalence of BD is probably less than 2% (1, 2), so the impact of undetected cases on power should be small (28). Another limitation arises from the fact that cases enrolled by different studies were ascertained differently, assessed with different diagnostic tools, and fulfilled different, albeit similar, diagnostic criteria. In any case, heterogeneity of ascertainment and clinical diagnosis would tend to reduce power, not lead to false positives (29, 30). This study included more than 2000 BD cases and twenty thousand controls that have not been included in previous GWAS of BD, but most of the cases analyzed in this study have been included in previous studies. Thus, the nominally significant support we observed for many loci identified in previous GWAS cannot be considered as independent evidence of replication. The purpose of this study was not to replicate previous findings but to identify additional risk loci for BD.

This study identified two novel loci associated with BD at genome-wide significance. The top SNP within the novel BD risk locus on chromosome 17q12 lies within an intron of *ERBB2* (Erb-B2 receptor tyrosine kinase 2). Data available through the GTEx Portal (http://www.gtexportal.org/) (31) suggests that this SNP acts as an expression quantitative trait locus (eQTL) for *ERBB2* in neural tissue (*P* = 2.6 × 10^−8^), but other eQTLs are also present at this locus. *ERBB2* is expressed in the brain and encodes a member of the epidermal growth factor (EGF) family of receptor tyrosine kinases, which can form homo-or hetero-dimers with other ERBB proteins. Interestingly, the ERBB proteins act as cell surface receptors for neuregulins (32). This pathway has long been thought to contribute to the pathogenesis of both BD and SCZ (33–37). Two previous gene expression studies have implicated *ERBB2* in BD (38, 39), but to our knowledge the present study is the first to demonstrate genetic association with BD. If confirmed in future studies, this locus might be a promising target for novel therapeutics.

The novel risk locus identified on chromosome 9p21.3 is within an intergenic region with no known protein coding genes nearby. A total of 17 highly linked variants in this region (spanning about 17 kb) met the genome-wide significance threshold. Genomic sequence alignments from multiple species suggest that the BD-associated segment is conserved in higher primates. The top SNP is in moderate LD with another SNP that lies within the promoter flanking region of *ELAVL2* which encodes a neuron-specific RNA binding protein that promotes neuronal development (40). Different SNPs at this locus have been previously implicated in a GWAS of SCZ (41).

Two previous GWAS have reported suggestive evidence of association between BD and common risk variants near *MAD1L1* (8, 26). The most significant SNP (rs4332037) reported in the PGC-BD (8) study, which was not genome-wide significant, lies about 54 kb away from --and is in low linkage disequilibrium (CEU r^2^ = 0.09) with -- our top SNP. Ruderfer and colleagues (26) identified *MAD1L1* as a genome-wide significant locus in a GWAS that used both BD and SCZ as a combined case definition. No previous study has to our knowledge demonstrated genome-wide significant association between this locus and BD only.

The MooDS-PGC study (9), which overlaps partially with the current study, identified two novel risk loci for BD. One of them is located on 6q16.1, a region between *MIR2113* and *POU3F2*. Our top SNP in this locus (rs1487441) is in perfect LD with rs12202969, the most significant variant in the original MooDS-PGC study. Thus the present study supports the MooDS-PGC finding in a larger sample, but cannot be considered an independent replication.

Another previously reported locus that is supported by the present study is on chromosome 12q13.1. Green et al. (24) reported that an intergenic polymorphism (rs7296288) between *RHEBL1* and *DHH* was significantly associated with BD. In the present study, an SNP (rs105442) in moderate LD (r^2^ = 0.62) with that reported by Green et al. was also significantly associated with BD. This SNP (rs105442) is located in the 3’–UTR of *DDN*, encoding dendrin. Dendrin is a synaptic protein that is markedly depleted by sleep deprivation (42), a common trigger of mood episodes in BD. Data available through the GTEx Portal (http://www.gtexportal.org/) (31) suggests that rs105442 acts as an eQTL locus for *DDN* in skeletal muscle tissue (*P* = 1.1 x 10^−19^).

In this study, we also carried out an X-chromosome meta-analysis that took advantage of the latest imputation methods and the best available analysis techniques (22, 43). The available sample size was underpowered to detect a common allele that confers a GRR as low as 1.2. Larger studies are needed to rule out the involvement of common variants within this range of GRR. However, the genomic heritability analyses showed that markers on the X-chromosome contributed very little to the total genomic heritability of BD. While the SNP density on this chromosome is lower than on the autosomes, the results suggest that there may be few, if any, X-linked markers that play an important role in risk for BD. The association method used in this study is powerful under the assumption of random X-inactivation (43), but may not be optimal when X-inactivation is skewed or incomplete (44). Some candidate-gene association studies (45, 46) have reported associations between BD and genes within the pseudoautosomal regions (PARs) of chromosome X. We cannot evaluate association with the PAR, since too few markers passed QC for inclusion in the present study.

This meta-analysis study has identified two novel risk loci for BD. The findings support previous work and also suggest novel genetic influences in BD. Increasing sample sizes should enable the identification of additional risk loci for BD, but X-linked markers seem to play a smaller than expected role in this common and severe mental illness.

## MATERIALS AND METHODS

### Overall study design

A detailed description of the study design and phenotype assessments for all samples can be found in the Supplementary Materials. This study was carried out in two-stages. Stage 1 comprised a meta-analysis of directly genotyped and imputed SNPs in 7,467 patients diagnosed with BD by direct interview and 27,303 controls drawn from published BD case-control studies, dbGaP, and WTCCC2. In Stage 2, a set of 179 variants from Stage 1 with *P* < 1.0 × 10^−6^ were tested in an independent sample of 2,313 BD cases and 3,486 controls from two ongoing Swedish GWAS.

### Stage 1 samples

Stage 1 comprised cases and controls from five published BD GWAS studies (Table 1) and an independent set of 2,072 European-ancestry patients with a long-term diagnosis of BD who were treated with lithium and submitted to the Consortium on Lithium Genetics (ConLiGen) (12). The ConLiGen sample has not been used in any previous GWAS studies of BD. An additional 20,998 genotyped individuals obtained from dbGaP and the WTCCC2 were used as controls. All participants included in the final association tests were of European ancestry.

### Stage 2 samples

The Stage 2 sample was predominantly Swedish and exclusively Northern European. BD cases from Sweden were collected through two recruitment streams: 1,908 cases from the Stanley study (47, 48) and 229 cases from the St. Göran Bipolar Project (49). Most control subjects (n=3,113) were randomly selected from Swedish population registers (ascertained on a national basis) and 55 were from the St. Göran Bipolar Project (49). The exclusion criterion for controls was any hospitalization for SCZ or BD. DNA collection procedures have been previously described (50). Both projects were approved by the Regional Ethical Review Board in Stockholm (Sweden), and all participants provided written informed consent. Genotyping was conducted at the Broad Institute of Harvard and MIT using Affymetrix 6.0 (Swedish sample 1) and Illumina OmniExpress (Swedish sample 2) chips.

### Quality control

Quality control procedures were carried out separately in each data set. The quality control parameters for retaining SNPs and subjects were: Subject missingness < 0.03; autosomal heterozygosity rate within mean ± 3SD; minor allele frequency (MAF) >= 0.01 (for Affymetrix data, we kept all variants with a MAF >= 0.05); SNP missingness < 0.05; and SNP Hardy-Weinberg equilibrium *P* > 10^−6^ in controls (for markers on the X chromosome, only females were used for the Hardy-Weinberg equilibrium tests), no discrepancies between reported sex and sex determined by genotypes on chromosome X. For each data set, around 100K LD-pruned SNPs were used to identify duplicated samples, cryptically related subjects, and population outliers. Relatedness testing was carried out by PLINK. Duplicated samples and cryptically related pairs were identified (Pi_hat > 0.10); only one member of each pair was kept for the data analyses. To identify cryptically related subjects present across different data sets, we tested approximately 20K LD-pruned SNPs present in all of the SNP arrays used by any of the studies. EIGENSOFT (51) was used to identify population outliers (Figure S5). All subjects with apparent non-European ancestry were excluded from the data analyses. After basic QC, we matched data sets by genotyping platform (Table 1).

### Imputation

Genotype imputation was performed using the prephasing/imputation strategy (52) by SHAPEIT2 (53) and IMPUTE2 (54) for both autosomes and the X chromosome. Phase 3 of the 1000 Genomes Project data set (n=2,504) was used as the reference panel for imputation. Appropriate options were used for X chromosome phasing and imputation (--chrX was set for SHAPEIT2 for phasing, and --chrX and gender information were provided to IMPUTE2 for imputation).

### Genotype-phenotype association analysis

Gene dosages for all markers with an imputation INFO ≥ 0.5 were used for the final association tests. In total, over 9M genotyped or imputed autosomal SNPs were analyzed. Genotype-phenotype association with imputed allele dosages for autosomal SNPs was carried out with logistic regression as implemented by PLINK (55). The first 10 principal components of population structure were used as covariates in the analyses.

Association tests for markers on the X chromosome were performed with Clayton's “one degree-of-freedom” test, implemented in the snpStats R package (22, 56). An extensive simulation study (43) of several different tests designed specifically for chromosome X association testing concluded that Clayton's “one degree-of-freedom” statistic is robust and powerful across a wide range of realistic conditions.

### Meta-analysis of Stage 1 samples

Meta-analysis in Stage 1 was conducted using the sample size-based method in METAL (57). Meta-analysis results were corrected with genomic control to eliminate any residual bias.

### Selection of SNPs for Stage 2

All markers with a p-value ≤ 1.0 × 10^−6^ in Stage 1 (n=179) were selected for validation in the Stage 2 samples, using identical QC and genotype-phenotype association procedures. Of these SNPs, results were available in the Stage 2 samples for 144 SNPs.

### Meta-Analysis of Stage 1 and Stage 2 samples

Meta-analysis of the Stage 1 and 2 results was conducted using the sample size-based method in METAL, under a fixed effects model. Again, genomic control was used to eliminate any residual bias. Association results were considered genome-wide significant if the meta p-values were < 5 × 10^−8^ and the heterogeneity tests were not significant (p>0.05).

### Identification of distinct association signals within risk loci

GCTA (58) was used to identify independently associated variants within each of the 6 GWAS-significant loci. GCTA made use of (i) summary statistics from the meta-analysis of the stage 1 and stage 2 samples, and (ii) genotype data from a reference sample for LD estimation between markers. Study-5 (416 BP cases and 2,741 controls) was used as the reference sample here because subjects from Study-5 were genotyped by the highest density SNP array (Illumina HumanOmni2.5M). GCTA uses a stepwise selection strategy (59) to identify any independent signals through an approximate conditional association analysis.

### Genomic heritability estimation of Stage 1 samples

We estimated the genomic heritability (the variance in case-control status explained by all genotyped markers) using a linear mixed model developed by Lee et al. (60). Briefly, a genome relationship matrix (GRM) between all pairs of individuals was estimated from the genome-wide markers. The GRM was then used to estimate variance attributable to markers via residual maximum likelihood (REML) analysis with the first 10 principal components of population structure used as covariates. The results were transformed to the liability scale by assuming a disease prevalence of 2% for BD. For the genomic heritability estimation, we only retained autosomal markers with MAF >0.01 and imputation INFO ≥ 0.9.

We partitioned genomic heritability attributable to each autosome by estimating the GRM from SNPs on each autosome and then fitting the GRMs of all the autosomes simultaneously in the linear mixed model. We further estimated the variance attributable to the X chromosome using a method developed by Yang and colleagues (58) under the assumption of full dosage compensation (complete inactivation of one X chromosome for females).

### Power estimation

Power analysis of autosomal markers was done with the Genetic Power Calculator (GPC) (61) under the following assumptions: Trait prevalence 2%, risk allele and marker allele frequency 25%, D-prime 1, GRR of 1.15 under a log-additive model. Since excess controls were used in this study and case/control ratio varied across sub-studies, simply using the total sample size might overestimate the power. To take this into account, we used GPC to estimate the noncentrality parameter (NCP) for each study and then iteratively determined the effective symmetric case/control sample size that returns the same NCP (62). The total effective sample sizes of the Stage 1 studies and Stage 1+2 studies were then used for the power estimation. Power analysis for markers on chromosome X was done with XGWAS (https://github.com/PeteHaitch/XGWAS), which uses a simulation-based method to estimate power. The total effective sample sizes of Stage 1 studies and Stage 1+2 were used for simulations.

## ACKNOWLEDGEMENTS

This work utilized the computational resources of the NIH HPC Biowulf cluster (http://hpc.nih.gov)

We obtained several datasets from the database of Genotypes and Phenotypes. A list of investigators and the funding support for these datasets can be found by searching http://www.ncbi.nlm.nih.gov/gap using these accession numbers: phs000124.v2.p1, phs000404.v1.p1, phs000303.v1.p1, phs000203.v1.p1, phs000170.v1.p1, phs000188.v1.p1, and phs000237.v1.p1

This study makes use of data generated by the Wellcome Trust Case-Control Consortium 2 (EGAD00000000021 and EGAD00000000023). We thank the Wellcome Trust Case Control Consortium for making data available for further analysis.

Some data and biomaterials were provided by the Rutgers Cell and DNA Repository.

Some data and biomaterials were collected in four projects that participated in the National Institute of Mental Health (NIMH) Bipolar Disorder Genetics Initiative. From 1991–98, the Principal Investigators and Co-Investigators were: Indiana University, Indianapolis, IN, U01 MH46282, John Nurnberger, M.D., Ph.D., Marvin Miller, M.D., Howard J. Edenberg, Ph.D. and Elizabeth Bowman, M.D.; Washington University, St. Louis, MO, U01 MH46280, Theodore Reich, M.D., Allison Goate, Ph.D., and John Rice, Ph.D.; Johns Hopkins University, Baltimore, MD, U01 MH46274, J. Raymond DePaulo, Jr., M.D., Sylvia Simpson, M.D., MPH, and Colin Stine, Ph.D.; NIMH Intramural Research Program, Clinical Neurogenetics Branch, Bethesda, MD, Elliot Gershon, M.D., Diane Kazuba, B.A., and Elizabeth Maxwell, M.S.W.

NIMH Study 1 -- Data and biomaterials were collected as part of ten projects that participated in the National Institute of Mental Health (NIMH) Bipolar Disorder Genetics Initiative. From 1999-03, the Principal Investigators and Co-Investigators were: Indiana University, Indianapolis, IN, R01 MH59545, John Nurnberger, M.D., Ph.D., Marvin J. Miller, M.D., Elizabeth S. Bowman, M.D., N. Leela Rau, M.D., P. Ryan Moe, M.D., Nalini Samavedy, M.D., Rif El-Mallakh, M.D. (at University of Louisville), Husseini Manji, M.D. (at Wayne State University), Debra A. Glitz, M.D. (at Wayne State University), Eric T. Meyer, M.S., Carrie Smiley, R.N., Tatiana Foroud, Ph.D., Leah Flury, M.S., Danielle M. Dick, Ph.D., Howard J. Edenberg, Ph.D.; Washington University, St. Louis, MO, R01 MH059534, John Rice, Ph.D, Theodore Reich, M.D., Allison Goate, Ph.D., Laura Bierut, M.D.; Johns Hopkins University, Baltimore, MD, R01 MH59533, Melvin McInnis, M.D., J. Raymond DePaulo, Jr., M.D., Dean F. MacKinnon, M.D., Francis M. Mondimore, M.D., James B. Potash, M.D., Peter P. Zandi, Ph.D, Dimitrios Avramopoulos, and Jennifer Payne; University of Pennsylvania, PA, R01 MH59553, Wade Berrettini, M.D., Ph.D.; University of California at Irvine, CA, R01 MH60068, William Byerley, M.D., and Mark Vawter, M.D.; University of Iowa, IA, R01 MH059548, William Coryell, M.D., and Raymond Crowe, M.D.; University of Chicago, IL, R01 MH59535, Elliot Gershon, M.D., Judith Badner, Ph.D., Francis McMahon, M.D., Chunyu Liu, Ph.D., Alan Sanders, M.D., Maria Caserta, Steven Dinwiddie, M.D., Tu Nguyen, Donna Harakal; University of California at San Diego, CA, R01 MH59567, John Kelsoe, M.D., Rebecca McKinney, B.A.; Rush University, IL, R01 MH059556, William Scheftner, M.D., Howard M. Kravitz, D.O., M.P.H., Diana Marta, B.S., Annette Vaughn-Brown, M.S.N., R.N., and Laurie Bederow, M.A.; NIMH Intramural Research Program, Bethesda, MD, 1Z01MH002810-01, Francis J. McMahon, M.D., Layla Kassem, PsyD, Sevilla Detera-Wadleigh, Ph.D, Lisa Austin, Ph.D, Dennis L. Murphy, M.D.

NIMH Study 19 -- Data and biomaterials were collected for the Systematic Treatment Enhancement Program for Bipolar Disorder (STEP-BD), a multi-center, longitudinal (5-8 year) project selected from responses to RFP #NIMH-98-DS-0001, “Treatment for Bipolar Disorder.” The project was led by Gary Sachs, M.D., and coordinated by Massachusetts General Hospital in Boston, MA. The NIMH grant number was 2N01MH080001-001.

NIMH Study 40 -- Data and biomaterials were collected as part of eleven projects (Study 40) that participated in the National Institute of Mental Health (NIMH) Bipolar Disorder Genetics Initiative. From 2003-2007, the Principal Investigators and Co-Investigators were: Indiana University, Indianapolis, IN, R01 MH59545, John Nurnberger, M.D., Ph.D., Marvin J. Miller, M.D., Elizabeth S. Bowman, M.D., N. Leela Rau, M.D., P. Ryan Moe, M.D., Nalini Samavedy, M.D., Rif El-Mallakh, M.D. (at University of Louisville), Husseini Manji, M.D. (at Johnson and Johnson), Debra A. Glitz, M.D. (at Wayne State University), Eric T. Meyer, Ph.D., M.S. (at Oxford University, UK), Carrie Smiley, R.N., Tatiana Foroud, Ph.D., Leah Flury, M.S., Danielle M. Dick, Ph.D (at Virginia Commonwealth University), Howard J. Edenberg, Ph.D.; Washington University, St. Louis, MO, R01 MH059534, John Rice, Ph.D, Theodore Reich, M.D., Allison Goate, Ph.D., Laura Bierut, M.D. K02 DA21237; Johns Hopkins University, Baltimore, M.D., R01 MH59533, Melvin McInnis, M.D., J. Raymond DePaulo, Jr., M.D., Dean F. MacKinnon, M.D., Francis M. Mondimore, M.D., James B. Potash, M.D., Peter P. Zandi, Ph.D, Dimitrios Avramopoulos, and Jennifer Payne; University of Pennsylvania, PA, R01 MH59553, Wade Berrettini, M.D., Ph.D.; University of California at San Francisco, CA, R01 MH60068, William Byerley, M.D., and Sophia Vinogradov, M.D.; University of Iowa, IA, R01 MH059548, William Coryell, M.D., and Raymond Crowe, M.D.; University of Chicago, IL, R01 MH59535, Elliot Gershon, M.D., Judith Badner, Ph.D., Francis McMahon, M.D., Chunyu Liu, Ph.D., Alan Sanders, M.D., Maria Caserta, Steven Dinwiddie, M.D., Tu Nguyen, Donna Harakal; University of California at San Diego, CA, R01 MH59567, John Kelsoe, M.D., Rebecca McKinney, B.A.; Rush University, IL, R01 MH059556, William Scheftner, M.D.,

Howard M. Kravitz, D.O., M.P.H., Diana Marta, B.S., Annette Vaughn-Brown, M.S.N., R.N., and Laurie Bederow, M.A.; NIMH Intramural Research Program, Bethesda, MD, 1Z01MH002810-01, Francis J. McMahon, M.D., Layla Kassem, Psy.D., Sevilla Detera-Wadleigh, Ph.D, Lisa Austin, Ph.D, Dennis L. Murphy, M.D.; Howard University, William B. Lawson, M.D., Ph.D., Evarista Nwulia, M.D., and Maria Hipolito, M.D. This work was supported by the NIH grants P50CA89392 from the National Cancer Institute and 5K02DA021237 from the National Institute of Drug Abuse.

The Swedish samples were funded by the Stanley Center for Psychiatric Research, Broad Institute from a grant from Stanley Medical Research Institute, the Swedish Research Council (K2014–62X–21445–05–3, K2012–63X–21445–03–2, K2014–62X–14647–12–51 and K2010–61P– 21568–01–4), the Söderström-Königska Foundation (SLS-472751), the Swedish foundation for Strategic Research (KF10-0039), and the China Scholarship Council. We would like to thank all of the participants who have kindly given their time and DNA to participate in our research, the Swedish quality register for bipolar disorders (BipoläR), and data collectors at the Department of Medical Epidemiology and Biostatistics (MEB) at Karolinska Institutet for help with recruitment of participants.

The ConLiGen project was in part funded by the Deutsche Forschungsgemeinschaft (DFG; grant no. RI 908/7–1) and the Intramural Research Program of the National Institute of Mental Health (ZIA-MH00284311; NCT00001174). The Romanian sample included in the ConLiGen project was also funded by UEFISCDI, Romania (grant no. 89/2012 awarded to Maria Grigoroiu-Serbanescu, PhD) and BMBF, Germany, grant no. 01GS08144 awarded to M. M. Nöthen and S. Cichon). The Canadian sample has been part of a study funded by the Canadian Institutes of Health Research (grant 64410 to MA). The collection of the Barcelona sample was supported by the Centro de Investigatión en Red de Salud Mental (CIBERSAM), IDIBAPS (grant numbers PI080247, PI1200906, PI12/00018) and Secretaria d’Universitats i Recerca del Departament d’Economia i Coneixement (2014SGR1636 and 2014SGR398).

Genotyping was funded in part by the German Federal Ministry of Education and Research (BMBF) through the Integrated Network IntegraMent (Integrated Understanding of Causes and Mechanisms in Mental Disorders), under the auspices of the e:Med Programme (grant 01ZX1314G to awarded to Thomas G. Schulze and Marcella Rietschel, grant 01ZX1314A to Sven Cichon and Markus M. Nöthen).

Thomas G. Schulze and Urs Heilbronner received support from the Dr.-Lisa-Oehler-Foundation (Kassel, Germany). Andrea Pfennig, Thomas Stamm, Michael Bauer, Andreas Reif, and Thomas G. Schulze received support from the German Federal Ministry of Education and Research (BMBF) within the framework of the *BipolLife* network (www.bipolife.org).

M.M.N. is a member of the DFG-funded Excellence-Cluster ImmunoSensation. M.M.N. also received support from the Alfried Krupp von Bohlen und Halbach-Stiftung. The study was supported by the German Research Foundation (DFG; grant ConLiGen RI 908/7– grant SFB 636 Z4; grant FOR2107, RI 908/11–1 to M.R., WI 3439/3–1 to S.H.W., NO 246/10–1 to M.M.N.).

The Genotype-Tissue Expression (GTEx) Project was supported by the Common Fund of the Office of the Director of the National Institutes of Health. Additional funds were provided by the NCI, NHGRI, NHLBI, NIDA, NIMH, and NINDS. Donors were enrolled at Biospecimen Source Sites funded by NCI\SAIC-Frederick, Inc. (SAIC-F) subcontracts to the National Disease Research Interchange (10XS170), Roswell Park Cancer Institute (10XS171), and Science Care, Inc. (X10S172). The Laboratory, Data Analysis, and Coordinating Center (LDACC) was funded through a contract (HHSN268201000029C) to The Broad Institute, Inc. Biorepository operations were funded through an SAIC-F subcontract to Van Andel Institute (10ST1035). Additional data repository and project management were provided by SAIC-F (HHSN261200800001E). The Brain Bank was supported by a supplements to University of Miami grants DA006227 & DA033684 and to contract N01MH000028. Statistical Methods development grants were made to the University of Geneva (MH090941 & MH101814), the University of Chicago (MH090951, MH090937, MH101820, MH101825), the University of North Carolina - Chapel Hill (MH090936 & MH101819), Harvard University (MH090948), Stanford University (MH101782), Washington University St Louis (MH101810), and the University of Pennsylvania (MH101822). The data used for the analyses described in this manuscript were obtained from the GTEx Portal on 10/12/2015.

The Australian cohort collection was supported by the Australian National Health and Medical Research Council (NHMRC) Program Grants 510135 (PBM) and 1037196 (PBM & PRS); and Project Grants 1063960 (JMF & PRS) and 1066177 (JMF).

The Swiss samples were funded by the Swiss National Science Foundation (grant number 32003B_125469 and NCCR Synapsy, Jean-Michel Aubry aand Alexandre Dayer).

Most importantly, we thank the individuals and families who have participated in and contributed time and data to these studies.

## ABBREVIATIONS

BD: Bipolar Disorder
SCZ: Schizophrenia
GWAS: Genome-wide Association Study
SNP: Single Nucleotide Polymorphism
MAF: Minor Allele Frequency
OR: Odds Ratio
SD: Standard Deviation
CI: Confidence Interval
eQTL: expression Quantitative Trait Locus
GAIN-NIMH: Genetic Association Information Network - National institute of Mental Health
WTCCC: Wellcome Trust Case Control Consortium
STEP-BD: Systematic Treatment Enhancement Program for Bipolar Disorder
TGEN: Translational Genomics Research Institute
ConLiGen: The International Consortium on Lithium Genetics
BoMA: Bonn-Mannheim
dbGaP: Database of Genotypes and Phenotypes
NCP: Non-Centrality Parameter
GRR: Genotype-Relative Risk

